# Circulating unacylated-ghrelin impairs hippocampal neurogenesis and memory in mice and is altered in human Parkinson’s disease dementia

**DOI:** 10.1101/259333

**Authors:** Amanda K. E. Hornsby, Vanessa V. Santos, Fionnuala Johnston, Luke D. Roberts, Romana Stark, Alex Reichenbach, Mario Siervo, Timothy Wells, Zane B. Andrews, David J. Burn, Jeffrey S. Davies

**Affiliations:** Molecular Neurobiology, Institute of Life Sciences, School of Medicine, Swansea University, UK.; Biomedical Discovery Institute, Department of Physiology, Monash University, Victoria, Australia.; Faculty of Medical Sciences, Newcastle University, UK.; School of Biosciences, Cardiff University, UK.

**Keywords:** Parkinson’s disease dementia, GOAT, acyl-ghrelin, unacylated-ghrelin, AG:UAG, adult hippocampal neurogenesis, memory

## Abstract

Blood-borne factors regulate adult hippocampal neurogenesis (AHN) and cognition in mammals, albeit via mechanisms that are poorly understood. We report that elevating circulating unacylated-ghrelin (UAG), using both pharmacological and genetic methods, reduced hippocampal neurogenesis and plasticity in mice. Spatial memory impairments observed in GOAT^-/-^ mice that lack acyl-ghrelin (AG) but have high levels of UAG, were rescued by treatment with AG. This unexpected finding suggests that the post-translational *acylation* of ghrelin is an important modulator of neurogenesis and memory in adult mammals. To determine whether this paradigm is relevant to humans we analysed circulating AG:UAG levels in Parkinsons disease (PD) patients diagnosed with dementia (PDD), cognitively intact PD patients and healthy controls. Uniquely, the ratio of plasma AG:UAG was reduced in the PDD cohort and correlated with cognitive performance. Our results identify UAG as a novel regulator of neurogenesis and cognition, and AG:UAG as a circulating diagnostic biomarker of dementia. The findings extend our understanding of adult brain plasticity regulation by circulating factors and suggest that manipulating the post-translational *acylation* of plasma ghrelin may offer therapeutic opportunities to ameliorate cognitive decline.

**Highlights:**

- Circulating UAG impairs markers of neurogenesis and plasticity
- GOAT^-/-^mice have impaired neurogenesis and spatial memory
- Circulating acyl-ghrelin (AG) rescues spatial memory deficit in GOAT^-/-^mice
- Circulating AG:UAG ratio is reduced in Parkinson’s disease dementia

## Introduction

Circulating factors are known to both enhance^1–4^ and impair^5–7^ neuronal plasticity and learning in adult mammals. However, the mechanisms underlying these effects are not completely understood. Systemic factors such as GDF11^1^ and TIMP2^4^ are reported to regulate the neural stem/progenitor cell (NSPC) niche in the adult hippocampus to promote new neurone formation and cognition. Conversely, circulating B2M^6^ and eotaxin^5^ impair the same niche resulting in reduced neurogenesis and impaired cognitive function. These data demonstrate that the hippocampal neurogenic niche is responsive to systemic factors, even in aged mammals, and suggest that circulating factors act as important modulators of mnemonic function. However, no single factor has been reported to fine-tune neurogenesis and cognition in both positive and negative directions. Here, we identify the post-translational acylation of the stomach hormone, ghrelin, as a novel mechanism controlling both up- and down-regulation of neurogenesis and learning in adult mice.

The birth and maturation of new neurones in the adult mammalian dentate gyrus (DG), termed adult hippocampal neurogenesis (AHN), is essential for spatial pattern separation memory^8,9^, which is the ability to separate highly similar components of memories into distinct memory representations^10^. This process is impaired in neurodegeneration^11^ and dementia^12^, but is enhanced by lifestyle factors, such as exercise^13^ and calorie restriction (CR)^14^. The molecular mechanisms governing proliferation and differentiation of NSPCs into new adult-born neurones are not fully understood and therapeutic strategies that promote AHN are limited.

We recently showed that CR increased AHN and hippocampal-dependent memory in a mechanism dependent on signaling via the stomach hormone, acyl-ghrelin^14^. Indeed, acyl-ghrelin, which is elevated during CR, crosses the blood-brain-barrier, binds to the growth hormone secretagogue receptor (GHSR) within the hippocampus and improves spatial memory^15^. Moreover, we showed that peripheral injection of acyl-ghrelin, at physiological doses, increases AHN and enhances pattern separation memory in adult rats^16^. Ghrelin must undergo post-translational acylation by the enzyme ghrelin-O-acyl transferase (GOAT)^17,18^, to form acyl-ghrelin, prior to binding and activating GHSR signaling^19^. Unacylated-ghrelin (UAG) represents ~80% of circulating ghrelin and is often considered an inactive precursor to acyl-ghrelin. Recent studies suggest that ghrelin may undergo tissue dependent acylation, including within the hippocampus, to support acyl-ghrelin signaling^20,21^. We therefore sought to determine whether UAG modulates AHN and hippocampal-dependent memory.

## Results and discussion

To assess whether UAG regulates adult NSPC plasticity in the sub-granular zone (SGZ) we analyzed the effect of peripherally administered UAG for seven-days in wild-type (WT) and GOAT-null (GOAT^-/-^) mice^17^. GOAT^-/-^ mice lack circulating acyl-ghrelin but have elevated levels of UAG making them ideally suited to assessing the loss of acyl-ghrelin coupled with increased plasma UAG^22^. Surprisingly, UAG-treated WT mice showed a significant 40% decrease in Ki67^+^ proliferating cells in the SGZ of the DG compared to vehicle-treated mice (Fig.1a,b). Similarly, genetic blockade of acyl-ghrelin signaling in GOAT^-/-^ mice reduced the number of dividing Ki67^+^ progenitor cells in the SGZ (Fig.1a,b). These findings were accompanied by a significant reduction in the number of doublecortin positive (Dcx^+^) immature neurons within the SGZ of both UAG-treated WT and GOAT^-/-^ mice (Fig.1c,d). Interestingly, UAG did not further exacerbate the reduction in proliferating cells and immature neurons in GOAT^-/-^ mice, suggesting that UAGs effect on neurogenesis may be via a saturable mechanism. While UAG is often considered inactive reports of UAG opposing acyl-ghrelin function on hepatocyte gluconeogenesis^23^, blocking acyl-ghrelin induced food intake^24^ and suppressing activity of the GH axis^25^, provide support for our findings. Together, these data suggest that the striking decrease in hippocampal cell proliferation and neurogenesis is due to elevated UAG, rather than simply loss of acyl-ghrelin.

**Figure 1.**
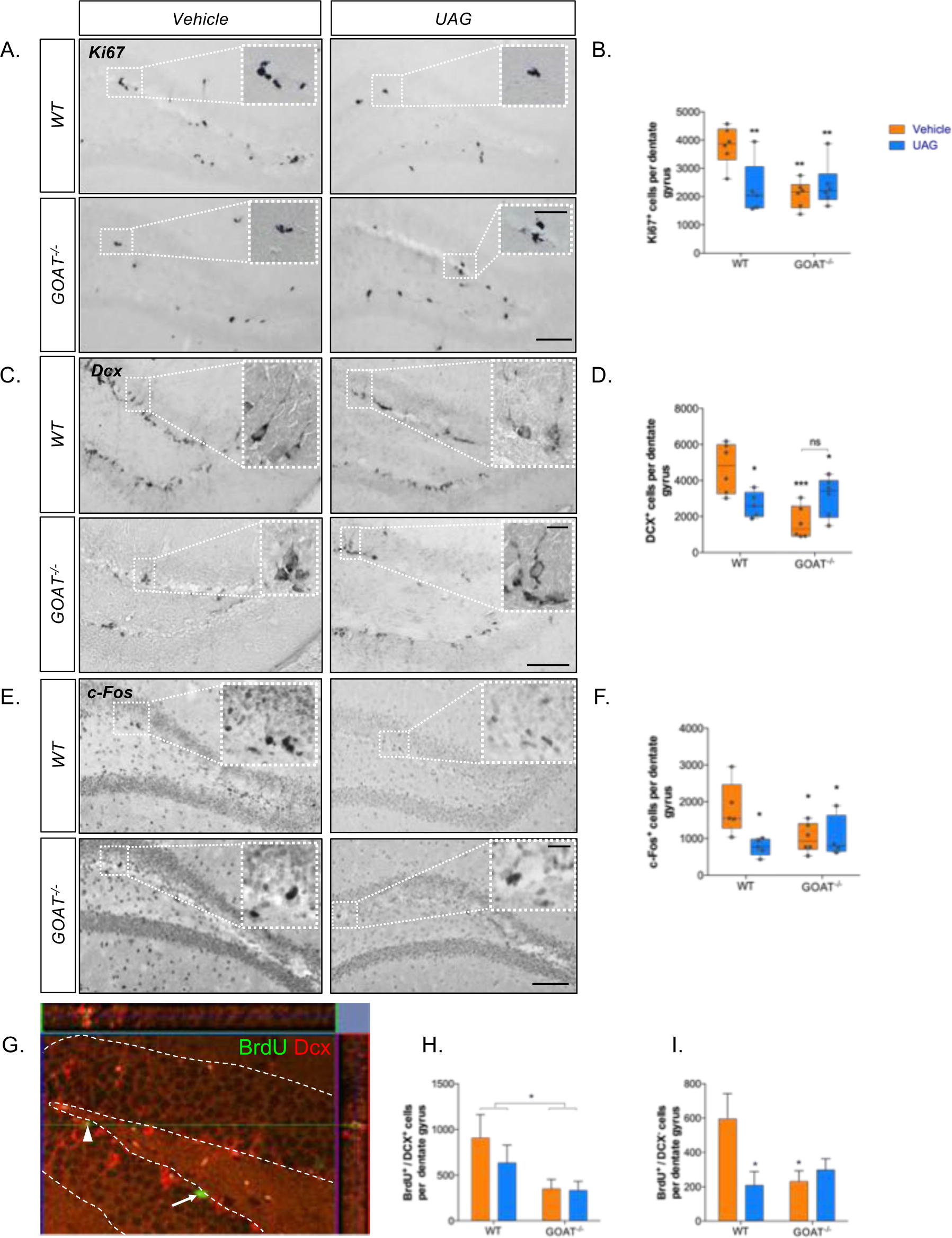
Unacylated-ghrelin (UAG) inhibits AHN in adult mice. Peripheral administration of UAG or genetic ablation of GOAT reduces the number of dividing Ki67^+^ cells (A,B], immature Dcx^+^ neurones (C,D] and c-Fos^+^ neurons (E,F] in the mouse DG. Representative confocal image showing new BrdU^+^/Dcx^+^ immature neurone (arrowhead] and a new BrdU^+^/Dcx” non-neuronal cell (arrow](G]. Genetic ablation of GOAT reduces survival of new neurons (main effect of genotype] (H] and new non-neuronal cells (I]. Statistical analysis was performed by 2-way AN OVA followed by Holm-Sidak *post-hoc* comparisons. Scale bar = 200nm (inset scale bar = 20 urn]. *P<0.05, **P<0.01, ***P<0.001 vs WT vehicle group. Data shown are mean ± SEM. *n =* 5–6 mice/group.

Next, we studied whether this UAG-mediated decrease in dividing progenitor SGZ cells reduced the survival of new adult-born DG neurons. To achieve this, mice in the above study received an injection of BrdU to birth-date proliferating cells on day 2 of the infusion prior to brain dissection and immunochemical analysis on day 7. The number of surviving immature neurons (BrdU^+^/Dcx^+^) was significantly reduced in GOAT^-/-^ mice (Fig.1g,h and Fig.S1b). Interestingly, we also demonstrated a significant reduction in the survival of new adult-born non-neuronal cells (BrdU^+^/Dcx^−^) in UAG-treated WT mice and in vehicle-treated GOAT^-/-^ mice, relative to vehicle-treated WT mice (Fig.1g,i).

Analysis of a second mouse model with genetic deletion of GOAT^18^ revealed a similar reduction in DG neurogenesis (Fig.S3a-d). Importantly, genetic deletion of GOAT (in either knockout model) did not affect the number of type II stem cells (Sox2^+^) in the SGZ (Fig.S1c,e and Fig.S3e,f), suggesting that the reduction in dividing progenitors was not due to a reduced stem cell pool but by modulation of the adult neurogenic niche. In addition, circulating immune factors implicated in reduced AHN, including eotaxin, fractalkine, IL-6, IL-10, RANTES and TNF-α, were not altered in GOAT^-/-^ mice (Fig.S2). These data identify GOAT-mediated acylation of ghrelin as a novel and important regulator of AHN.

Our findings are notable as we show that ghrelin^-/-^ mice, which lack both acyl-ghrelin and UAG, do not have impairments in basal adult neurogenesis. Previously, the rate of cell proliferation and neuronal differentiation was reported as being reduced in ghrelin^-/-^mice^26^. However, our analyses of hippocampal NSPC number (Sox2), cell proliferation (Ki67), immature neuron number (Dcx) and new adult-born neuron number (BrdU/NeuN), show no change compared to wild-type mice (Fig.S4). These data from ghrelin^-/-^ mice are consistent with the loss of both the pro-neurogenic acyl-ghrelin and the anti-neurogenic UAG resulting in no net change in AHN. In contrast, GOAT^-/-^ mice lack acyl-ghrelin but have high levels of UAG, suggesting that high UAG is responsible for reduced AHN rather than the loss of acyl-ghrelin.

Next, we assessed whether elevation of UAG disrupted other molecular signatures of DG impairment. Indeed, we observed a significant reduction in immuno-labeling of the immediate early gene, c-Fos, in both UAG treated WT (58%) and GOAT^-/-^ (44%) mice (Fig.1e,f). Given the role of c-Fos in neural plasticity and memory consolidation^27–29^ we reasoned that hippocampal-dependent learning and memory may be impaired in these mice.

To determine whether the neurochemical deficits observed in GOAT^-/-^ mice resulted in memory impairments and whether this could be rescued by acyl-ghrelin, adult WT and GOAT^-/-^ mice were given daily injections of either saline or acyl-ghrelin for one or seven days prior to analysis of hippocampal-dependent spatial memory using a Y-maze (Fig.S3a-d). GOAT^-/-^ mice displayed a deficit in performance that was unaffected by injection of acyl-ghrelin 1h prior to testing (Fig.2a). However, treatment with acyl-ghrelin for seven days rescued the memory deficit in GOAT^-/-^ mice (Fig.2b). Interestingly, this enhancement in spatial memory performance remained partially intact when the mice were tested on day 28, 21 days following the end of treatment (Fig.2c). The long-term rescue of spatial memory, long after acyl-ghrelin has cleared the circulation, is consistent with acyl-ghrelin-mediated AHN16. Further work using behavioral tests that place emphasis on distinguishing similar but distinct spatial contexts is required to fully elucidate the consequences of reduced AHN in GOAT^-/-^ mice. However, these data are consistent with our neurochemical findings and demonstrate that GOAT is essential for hippocampal-dependent spatial memory.

**Figure 2.**
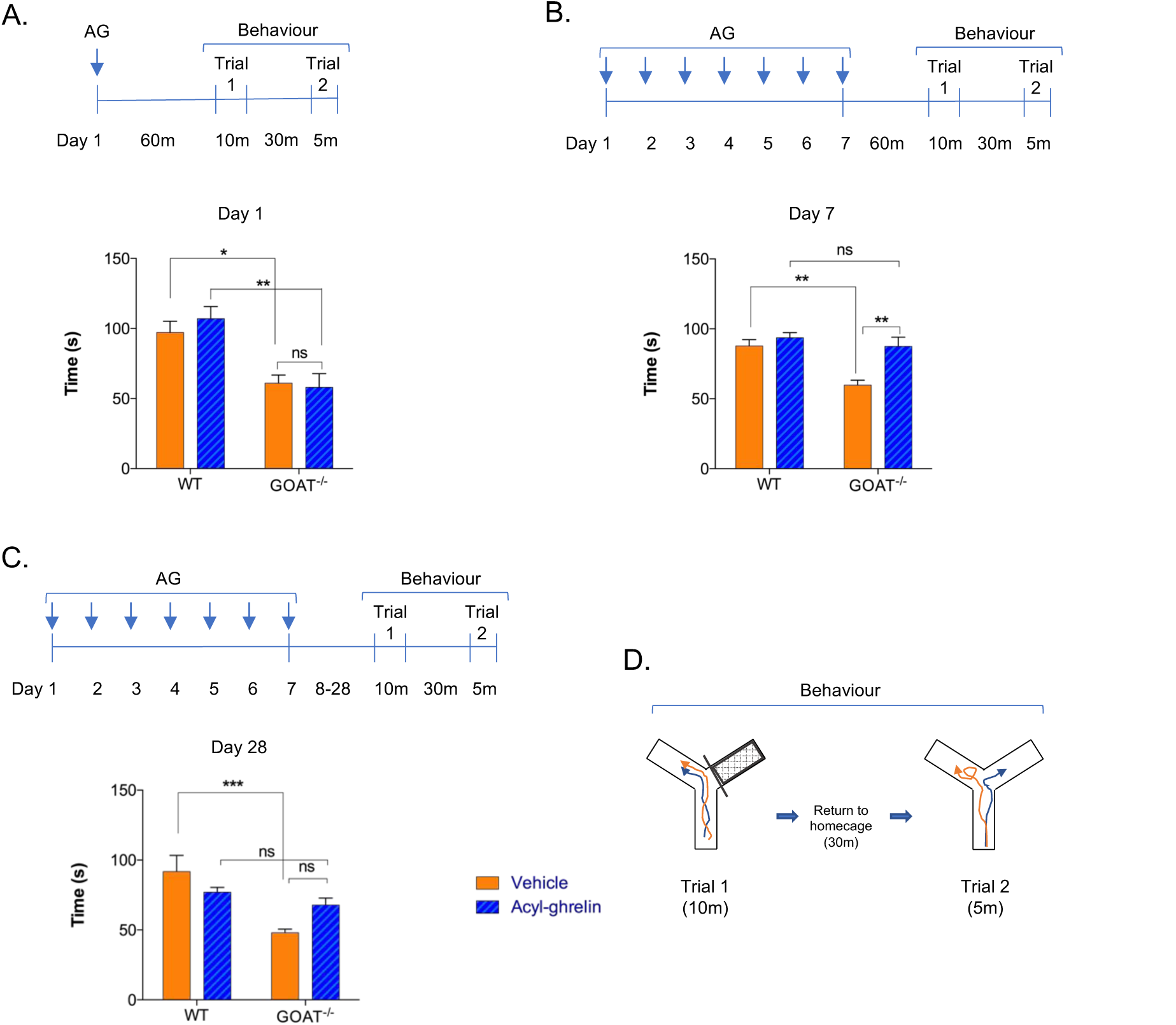
Adult GOAT/” mice display hippocampal-dependent spatial memory deficits that are rescued by acyl-ghrelin treatment. GOAT^-/-^ mice display memory impairments (A) that are restored by acyl-ghrelin treatment for 7 days (B) and 21 days following the end of treatment on day 28 (C). Schematic representations of treatment timelines (A-C) and behavior using Y-maze (D) are shown. Statistical analysis was performed by 2-way ANOVA followed by Tukey’s *post-hoc* comparisons *[n=6* mice/group). *P<0.05, **P<0.01, ***P<0.001. All data shown are mean ± SEM.

Mechanistically, we reasoned that as circulating levels of acyl-ghrelin and UAG have opposing actions on neurogenesis and cognition in mice, there should be a reduction in the plasma ratio of acyl-ghrelin to UAG (AG:UAG) in humans diagnosed with dementia. As acyl-ghrelin protects against neuron loss in models of Parkinson’s disease (PD)^30,31^ and is reportedly reduced in human PD^32,33^, we hypothesized that circulating AG:UAG ratios may be particularly affected in individuals diagnosed with PD dementia (PDD) compared to a cognitively-unimpaired PD group. To test this hypothesis, we recruited individuals with PD, PDD and age-matched healthy controls to determine fasting and post-prandial levels of both acyl-ghrelin and UAG. In keeping with our pre-clinical findings, we found that the plasma ratio of AG:UAG was significantly reduced in the PDD group compared to the cognitively intact PD and control cohorts (Fig.3a,b). Consistent with this finding, cognitive impairment was correlated with a reduction in plasma AG:UAG (Fig.3c), suggesting this ratio as a potential novel diagnostic biomarker for human dementia. Interestingly, the cognitively intact PD group did not show reduced acyl-ghrelin levels in either the fasted or 180 minutes after eating (Table S1), further supporting a specific role for ghrelin in regulating mnemonic function. To the best of our knowledge this is the first study of ghrelin in PD to perform stratification of patients by cognitive ability.

**Figure 3.**
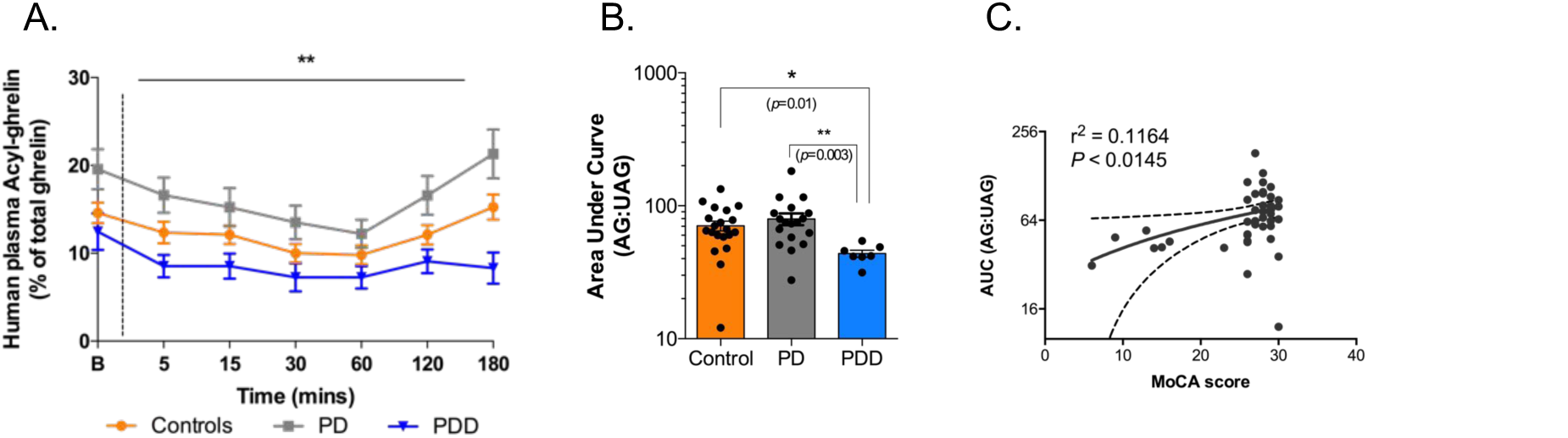
The ratio of AG:UAG is reduced in humans with PD dementia (PDD). Plasma AG:UAG ratio in healthy controls (n=20), PD (n=20) and PDD (n=8) patients under fasting and post-prandial conditions. Dotted line indicates breakfast consumption at time 0 (A). Area under curve (AUC) values demonstrate a significant reduction in AG:UAG ratio in control vs PDD and PD vs PDD groups (B). Correlation of cognition (MoCA score) with plasma AG:UAG (AUC) (C). Statistical analysis was performed by Kruskall-Wallis test with Dunns post-hoc multiple comparison and Spearman correlation analysis (*two-tailed*). *P<0.05, **P<0.01. All data shown are mean ± SEM.

Notably, there is impaired hippocampal plasticity, including neurogenesis, in rodent toxin-based and genetic-based models of PD^34–36^, whilst post-mortem human PDD brain have reduced numbers of NSPCs and immature neurones in the adult DG^11^. Moreover, PD patients display impaired performance in the Object Pattern Separation task which is consistent with impaired DG function and possibly reduced AHN^37^. Combined, these studies provide a putative mechanistic link between AG:UAG ratio, AHN and PDD in humans.

Together, our findings demonstrate a previously unknown function for UAG and GOAT in reducing neurogenesis and spatial memory performance, and suggest that enhancing circulating acyl-ghrelin, or more importantly, increasing the AG:UAG ratio may ameliorate cognitive decline. To the best of our knowledge this is the first description of a post-translational modification to a circulating factor that modulates, in either direction, neurogenesis and cognition.

## Methods

### Study approval

The animal procedures described, including those involving genetically modified animals, conformed to the U.K. Animals (Scientific Procedures) Act 1986 and the Monash University Animal Ethics Committee guidelines. All procedures complied with the NC3Rs ARRIVE Guidelines on reporting *in vivo* experiments. The study involving humans was approved by Local Ethical Review (14NE0002) at Newcastle University. A complete description of the methods is provided in Supplemental material.

### Animals

Six-month old homozygous GOAT-null (GOAT^-/-^) mice and their WT (C57Bl6) littermate controls17 were imported from Taconic Farms (Hudson, NY) and housed in the JBIOS animal facility (Cardiff University) under standard laboratory conditions (12h L:D cycle) with food and water available *ad libitum.* Six-month old Ghrelin-null (Ghrelin^-/-^) mice and their WT littermates^29^ were bred from heterozygous x heterozygous matings in the animal facility at Cardiff University under standard conditions (as above). Founder stock were a kind gift from Prof. Yuxiang Sun (Baylor College of Medicine, Houston, USA). A second genetic model of GOAT ablation^18^ was provided by Regeneron Pharmaceuticals. This GOAT^-/-^ line was generated using Velocigene technology. The GOAT gene sequence (ATG-stop) was replaced with a lacZ reporter gene using the target vector, bacterial artificial chromosome (BAC). These mice originated from C57BL/6/129 targeted embryonic stem cells and mice were backcrossed onto a C57BL/6 mice background. Mice were kept in standard laboratory conditions at Monash University with free access to food (chow diet, cat no. 8720610 Barastoc stockfeeds, Victoria Australia), and water at 23°C in a 12-hour light/dark cycle and were group-housed to prevent isolation stress, unless otherwise stated.

### UAG infusion

Each mouse^17^ was fitted with an indwelling jugular vein catheter connected to a subcutaneous osmotic mini-pump (ALZET model 1007D) primed to deliver vehicle (sterile isotonic saline containing BSA (1mg/ml) and heparin (5U/ml) at 0.5µl/h) or UAG (48ug/day) under isoflurane anesthesia. One day later all mice received an injection of thymidine analogue, BrdU (50mg/kg i.p), to label dividing cells. After seven days mice were re-anesthetized and killed by decapitation, the brains being excised whole and processed for immunohistochemistry (IHC) as described below.

### Tissue collection

Whole brain was removed and immediately fixed by immersion in 4% paraformaldehyde (PFA) in 0.1M phosphate buffer (pH 7.4) for 24h at 4°C. Subsequently brains were cryoprotected in 30% sucrose solution (until sunk). Coronal sections (30μm) were cut into a 1:6 series along the entire rostro-caudal extent of the hippocampus using a freezing-stage microtome (MicroM, ThermoScientific) and collected for IHC. All IHC was performed on free-floating sections at room temperature unless stated otherwise.

### Immunohistochemistry

For immunofluorescent analysis of Brdu^+^/Dcx^+^, sections were washed three times in PBS for 5 minutes, permeabilized in methanol at −20°C for 2 minutes and washed (as before) prior to pre-treatment with 2N HCl for 30 minutes at 37°C followed by washing in 0.1 M borate buffer, pH8.5, for 10 minutes. Sections were washed as before and blocked with 5% normal donkey serum (NDS) in PBS plus 0.1% triton (PBS-T) for 60 minutes at room temperature. Sections were incubated overnight at 4°C in rat anti-BrdU (1:400, AbD Serotec) and goat anti-Dcx (1:200, Santa Cruz Biotechnology, USA) diluted in PBS-T. Tissue were washed as before and incubated in donkey anti-rat AF-488 (1:500, Life Technologies, USA) and donkey anti-goat AF-568 (1:500, Life Technologies, USA) in PBS-T for 30 minutes in the dark. After another wash sections were mounted onto superfrost+ slides (VWR, France) with prolong-gold anti-fade solution (Life Technologies, USA). For immunofluorescent analysis of Sox2, sections were treated as above with the exception of antigen retrieval being performed in sodium citrate at 70°C for 1h (rather than 2N HCl or borate buffer) with subsequent blocking in 5% NGS. Immunoreactivity was detected using rabbit anti-Sox2 (1:500, ab97959, Abcam) and goat anti-rabbit AF-568 (Life Technologies, USA). Nuclei were counterstained with Hoechst prior to mounting as described above.

For DAB-immunohistochemical analysis of Ki67, Sox2 and DCX labeling, sections were washed in 0.1M PBS (2x10mins) and 0.1M PBS-T (1x10 mins). Subsequently, endogenous peroxidases were quenched by washing in a PBS plus 1.5% H2O2 solution for 20 minutes. Sections were washed again (as above) and incubated in 5% NGS (NDS for DCX) in PBS-T for 1h. Sections were incubated overnight at 4°C with rabbit anti-Ki67 (1:500, ab16667, Abcam), rabbit anti-Sox2 (1:1000, ab97959, Abcam) or goat anti-DCX (1:200 Santa Cruz Biotechnology, USA), in PBS-T and 2% NGS (NDS for DCX) solution. Another wash step followed prior to incubation with biotinylated goat anti-rabbit (1:400 Vectorlabs, USA) for Ki67 and Sox2 or biotinylated donkey anti-goat (1:400 Vectorlabs, USA) for DCX, in PBS-T for 70 minutes. The sections were washed and incubated in ABC (Vectorlabs, USA) solution for 90 minutes in the dark prior to another two washes in PBS, and incubation with 0.1M sodium acetate pH6 for 10 minutes. Immunoreactivity was developed in nickel-enhanced DAB solution followed by two washes in PBS. Sections were mounted onto superfrost+ slides (VWR, France) and allowed to dry overnight before being dehydrated and de-lipified in increasing concentrations of ethanol. Finally, sections were incubated in Histoclear (2x3 mins; National Diagnostics, USA) and coverslipped using entellan mounting medium (Merck, USA). Slides were allowed to dry overnight prior to imaging.

For DAB-immunohistochemical analysis of c-Fos labeling, sections were washed and endogenous peroxidases quenched as before. Sections were washed again (as above), before antigen retrieval in sodium citrate at 70°C for 1h and subsequent blocking in 5% NGS in PBS-T for 1h. Sections were incubated overnight at 4°C with goat anti-rabbit anti-c-Fos (1:4000, SC-52, Santa Cruz, USA) in PBS-T and 2% NGS solution. Another wash step followed prior to incubation with biotinylated goat anti-rabbit (1:400 Vectorlabs, USA) in PBS-T for 70 minutes. The sections were washed and incubated in ABC (Vectorlabs, USA) solution for 90 minutes in the dark prior to another round of washing (as above) and subsequent tyramide signal amplification. Following incubation with biotinylated tyramine (1:100) in PBS-T plus 0.1% H_2_O_2_ for 10 min, sections were washed in 0.1M PBS (1x10 mins) and 0.1M PBS-T (2x10 mins), before a second 90 minutes ABC (Vectorlabs, USA) incubation, which was again performed in the dark. Sections were then washed in 0.1M PBS (2x10 mins) and incubated with 0.1M sodium acetate pH6 for 10 minutes. Immunoreactivity and tissue processing was performed as described above.

### Quantification of labeled cells

Immuno-stained brain tissue was imaged by light microscopy (Nikon 50i) (for DAB), fluorescent (Axioscope, Zeiss) or confocal microscopy (LSM710 META, Zeiss). Immunolabelled cells were manually counted bilaterally through the z-axis using a ×40 objective and throughout the rostro-caudal extent of the granule cell layer (GCL). Resulting numbers were divided by the number of coronal sections analyzed and multiplied by the distance between each section to obtain an estimate of the number of cells per hippocampus (and divided by 2 to obtain the total per DG). For quantification of DG volume, Hoechst nuclear stain was used on tissue sections as above and fluorescent area expressed as μm^2^ per section. Images were processed using Zen (Zeiss) or Image J software. All analyses were performed blind to genotype and treatment.

### Milliplex plasma analysis

Mouse plasma was analyzed using the Milliplex-MAP 6-plex mouse cytokine magnetic bead panel kit (Cat #SPR402), to analyze the cytokines IL-6, eotaxin, fractakine, IL-10, RANTES and TNFα. The assay was performed according to the manufacturers guidelines. All reagents were brought to room temperature (RT) before use in the assay. Plasma samples were thawed at 4°C and diluted 1:2 with assay buffer. Antibody-immobilized beads were sonicated for 30 seconds and vortexed for 1 minute. 60µl from each antibody-bead vial was added to the mixing bottle provided and the final volume made up to 3ml with assay buffer. Wash buffer (WB) was prepared by mixing 60ml 10X WB with 540ml deionized water. Serum matrix solution was prepared by adding 2ml assay buffer to the lyophilized serum matrix. Subsequently, the mouse cytokine standard cocktail (Cat #MXM8070) was reconstituted with 0.25ml deionized water and a 1:5 serial dilution was made.

WB (200µl) was added to each well of the 96-well plate before it was sealed and agitated for 10 minutes at room temperature. The WB was removed and 25µl of each standard or control was added to the appropriate wells and 25µl of assay buffer to the sample wells. Next, 25µl of serum matrix was added to the background, standards and control wells, and 25µl of diluted plasma sample was added into the sample wells. Following this, the mixing bottle containing the antibody-bead mixture was vortexed and 25µl was added to each well. The plate was sealed, wrapped in foil and incubated at 2–8°C on a plate shaker for 16–18 hours.

After incubation, the well contents were removed (using a handheld magnet) and the wells washed twice with 200µl WB. Subsequently, 25µl of detection antibodies were added to each well, the plate was then sealed, covered with foil and incubated for 1h at room temperature, with agitation. Next, 25µl Streptavidin-Phycoerythrin was added to each well and incubated for 30 minutes as before. Finally, with a hand-held magnet, well contents were removed and the plate was washed twice with 200µl WB. 150µl of sheath fluid (BioRad Bio-Plex Sheath Fluid, Cat #171–000055) was added to each well for 5 minutes with agitation. The plate was assessed using a Bio-Rad Bioplex-200 System with Bio-Plex Manager 4.1 software.

### Behavioral testing

To determine whether the neurochemical deficits observed in GOAT^-/-^ mice resulted in impaired hippocampal-dependent spatial memory and whether this could be rescued by acyl-ghrelin, adult 12 week-old WT and GOAT^-/-^ mice (n=6/group) were given daily injections of either saline or acyl-ghrelin (300μg/kg i.p) for 1 or 7 days prior to analysis of spatial memory using a Y-maze. This dose of acyl-ghrelin was chosen as it has previously been shown to increase food intake. Injections were performed daily between 9-10am when mice were in a fed state. A modified Y-maze task was used to assess spatial memory performance on day 1, 7 or 28 days after the first acyl-ghrelin injection. All tests were performed in an experimental room with sound isolation and dim light. The animals were carried to the test room for at least 1 hour of acclimation. Behavior was monitored using a video camera positioned above the apparatuses and the videos were later analyzed by an experienced blinded researcher using video tracking software (CleverSys Inc, Reston, VA, USA). The modified Y-maze measures spatial memory, as spatial orientation cues facilitate rodents to explore a novel arm rather than returning to a previously visited arm. We used a Y-shaped grey Perspex maze (30 cm x 10 cm x 16 cm) and each arm could be isolated by blocking entry with a sliding door. Saw dust from a mouses home cage lined the maze during the trials and extra maze cues on the walls were placed 30–40 cm from the end of the arms to provide spatial orientation cues. Behavior was tested across two trials, the first of which had one arm of the maze randomly blocked off. Mice were allowed to explore the reduced maze for 10 minutes and then returned to their home cage. The second trial was conducted 30 minutes after the first trial and both arms of the maze were opened. Mice were placed in the start arm and allowed to explore the full maze for 5 minutes. All behaviors were recorded and analyzed using tracking software. Novel arm exploration was recorded when all four feet of each mouse entered the novel arm. The apparatus was cleaned with 80% ethanol between each trial and each animal.

### Human plasma collection

All procedures involving human participants were performed at the Clinical Ageing Research Unit, Newcastle University, with appropriate ethical approvals. 48 adults aged 60–85 were recruited; healthy controls (HC) (n=20), PD (n=20) and PDD according to level 1 Movement Disorder Task Force criteria for the diagnosis of PDD38 (n=8). Montreal Cognitive Assessment (MoCA) was ≥ 26/30 for HC and PD, and ≤25/30 for PDD. Participants with UWL, obesity, BMI <18 or >30, diabetes, gastrointestinal disease, smoking, deep brain stimulation or non-selective anticholinergic medication were excluded. Participants were tested fasted and off PD medication. Blood was drawn in the fasted state (0) and at 5,15, 30, 60,120 and 180 minutes following a standard breakfast. Blood samples for assessing acyl-ghrelin were treated with 4-(2-Aminoethyl)-benzenesulfonyl fluoride to prevent de-acylation and analyzed using a multiplex assay (Millipore). Total ghrelin was analyzed by ELISA (Millipore). Area under the curve (AUC) was calculated for each analyte. Three outliers were identified using the ROUT test (Q=1%) and removed prior to analysis using a Kruskall-Wallis test with Dunns post-hoc multiple comparison.

### Statistical analysis

Statistical analyses were carried out using GraphPad Prizm 6.0 for Mac. Data distribution were assessed using the Schapiro-Wilks normality test. For normally-distributed data, comparisons between two groups was assessed by two-tailed unpaired Students *t*-test. For multiple groups with one variable factor, a one-way ANOVA was used, and where there were two variable factors a two-way ANOVA was used. Appropriate post-hoc tests were used as described. Data displaying non-Gaussian distribution were analyzed by non-parametric tests, as described in the text. Spearman correlation *(two-tailed)* and linear regression analysis were used to determine the goodness-of-fit between plasma AG:UAG (AUC) and cognition (MoCA). Data are presented as mean ± sem. *, P < 0.05; **, P < 0.01; ***, P < 0.001 were considered significant.

## Author Contributions

AH, VV, FJ, LR, RS, AR, MS, TW performed the experiments and generated the data. TW, ZBA, DB and JSD designed the study. JSD prepared the figures and wrote the manuscript. All authors reviewed the manuscript.

## Acknowledgements

The authors would like to thank Dr Jesus A Gutierrez (Translational Science & Technologies, Eli Lilly & Co., Indianapolis, IN, USA) for the supply of GOAT^-/-^ mice. We would also like to thank Mrs Sally James (Swansea University) for assistance with confocal microscopy. This work was funded by grants from the Medical Research Council (Grant no. G0902250, to JSD), the St Davids Medical Foundation (to JSD), British Society for Neuroendocrinology (to ZBA and JSD), The Royal Society (to JSD) and the National Institute for Health Research Newcastle Biomedical Research Unit based at Newcastle Hospitals Foundation Trust and Newcastle University (to DB and JSD).

## Supplemental information

**Figure SI.**
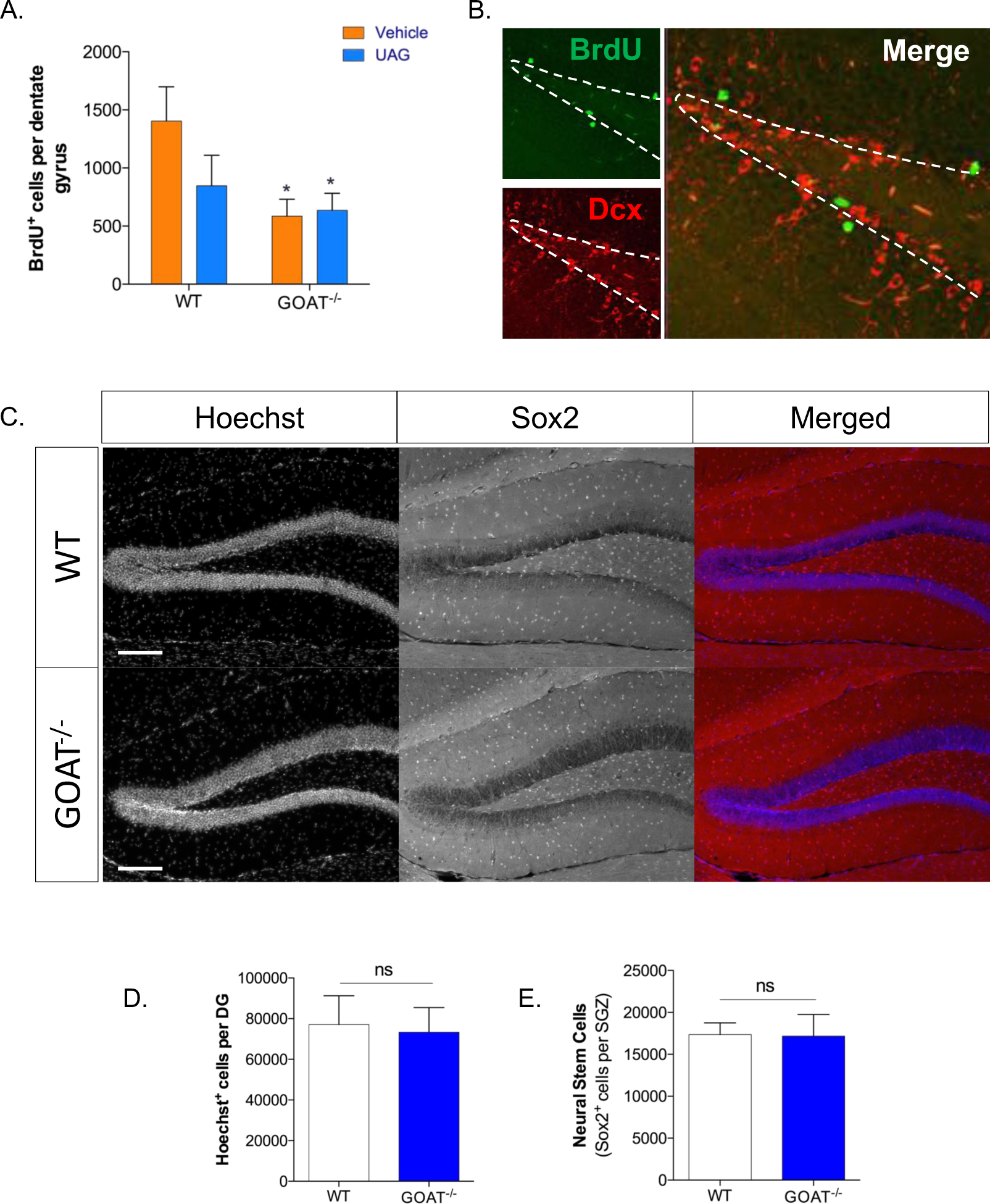
GOAT^-/-^ mice display reduced new BrdU^+^ cell number in the DG compared to WT mice [A]. Representative confocal image showing BrdU [green] and Dcx [red] in the mouse hippocampus [B]. Statistical analysis was performed by 2-way ANOVA followed by Holm-Sidak *post-hoc* comparisons vs WT vehicle group [n=5–6 mice/group]. Fluorescent images of Hoechst^+^ nuclei and Sox2^+^ NSPCs in DG from WT and GOAT/” mice [C]. GOAT/” mice do not have gross anatomical changes [Hoechst^+^][D] or impaired NSPC [Sox2^+^][E] cell number in the hippocampal DG. Statistical analysis performed by two-tailed Student’s *t-*test [n=4 mice/group]. *P<0.05. Scale bar = 200nm. All data shown are mean ± SEM.

**Figure S2.**
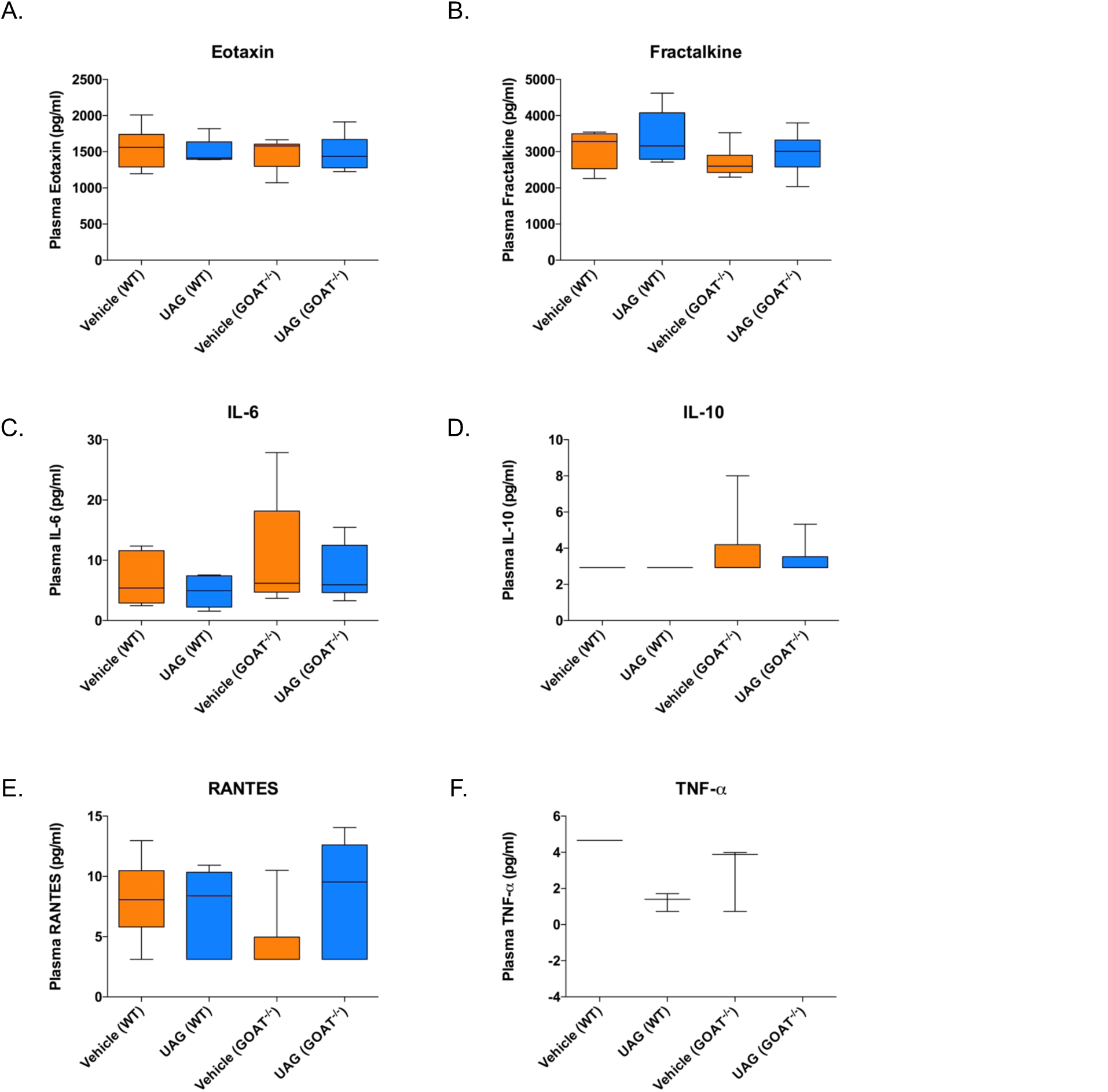
Peripheral administration of unacylated-ghrelin (UAG) or genetic ablation of GOAT does not affect quantities of circulating factors that are known to modulate neurogenesis (A-F). Statistical analysis was performed by 2-way ANOVA followed by Holm-Sidak *post-hoc* comparisons vs WT vehicle group. *P<0.05. All data shown are mean ± SEM. N = 5–6 mice/group.

**Figure S3.**
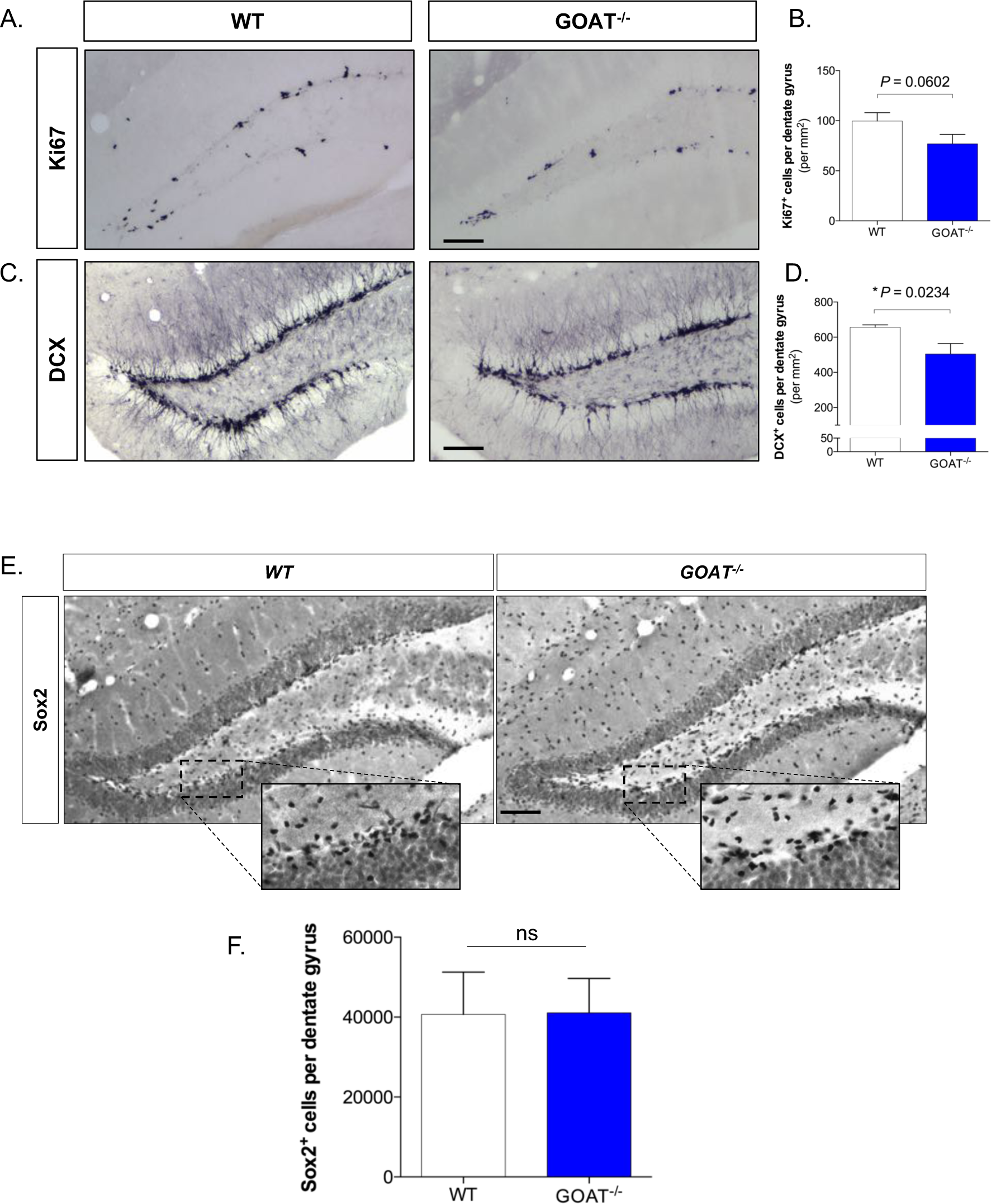
GOAT^-/-^mice, generated by Yang et al.^17^, have a reduced rate of cell proliferation (A,B) and neurogenesis (C,D) in the hippocampal DG. However, NSPC (Sox2^+^) number in the SGZ of the hippocampal DG is similar between WT and GOAT^-/-^mice (E,F). Statistical analysis was performed by two-tailed Students t-test. Scale bar = 200|mi. *P<0.05. All data shown are mean ± SEM. *n* = 6 mice/group.

**Figure S4.**
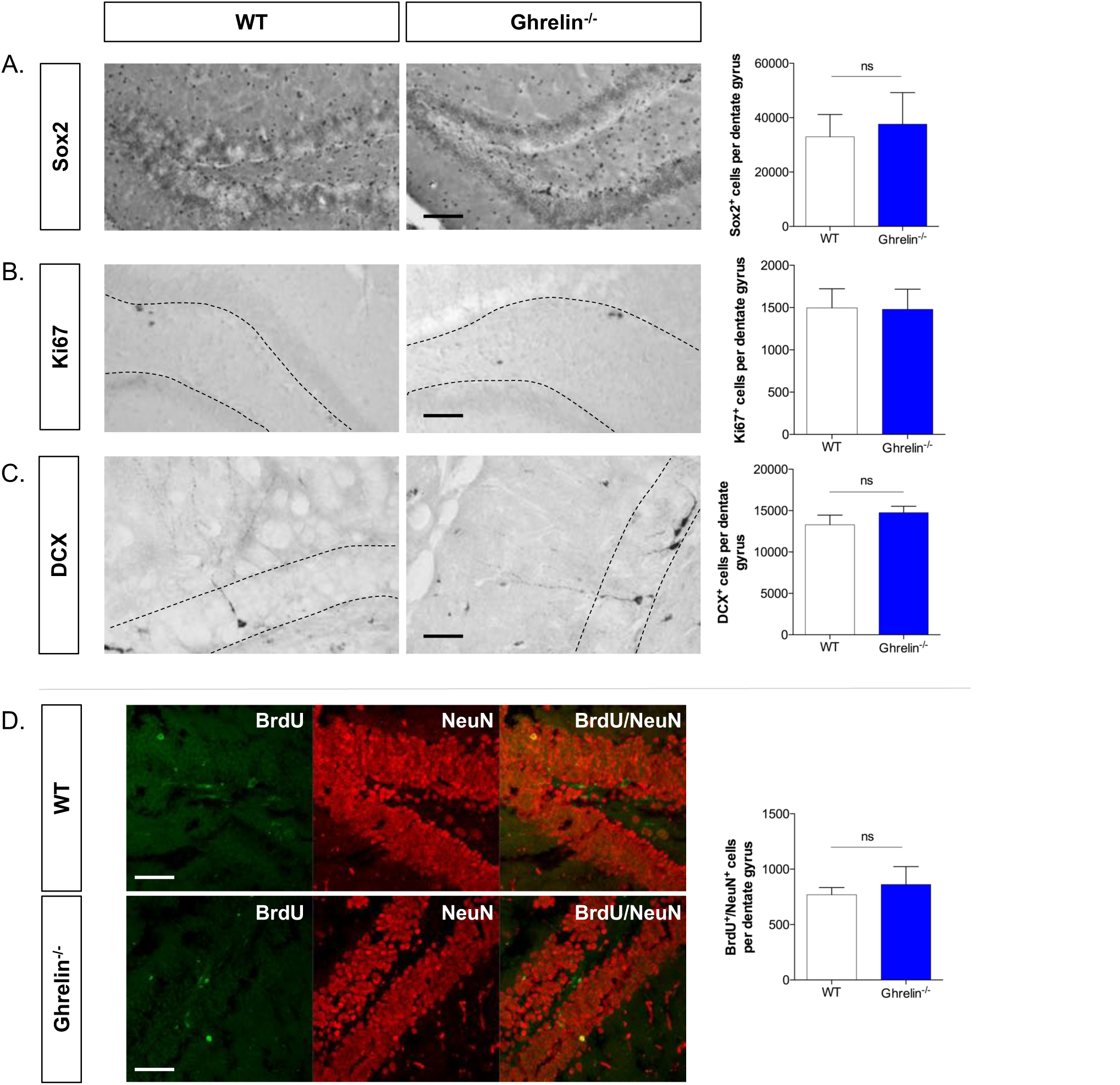
Ghrelin^-/-^ mice have unaltered numbers of Sox2^+^ neural stem cells (*n*=8/group) (A), Ki67^+^ dividing NSPCs (WT *n*=4/group)(B), Dcx^+^ immature neurons (WT *n=9*, ghrelin^-/-^*n=12)[C)* and BrdU^+^/NeuN^+^ new adult born neurons (WT *n=6*, ghrelin^-/-^ *n=7)* (D) in the hippocampal DG. Statistical analysis was performed by two-tailed Students t-test. Scale bar = 200|im for A&B, 50|im for C&D. ns = not significant. All data shown are mean ± SEM.

**Table S1.** Demographic information for study participants. MoCA; Montreal Cognitive Assessment. BMI; Body Mass Index. Statistical analysis was performed by Kruskal Wallis test or the Chi-squared test^+^. \**P* <0.05 considered significant.

|  | Healthy controls<br>(n=20) | Parkinson's disease<br>(n=20) | Parkinson's disease dementia<br>(n=8) | Significance |
| --- | --- | --- | --- | --- |
| Age<br>(mean ± sd) | 74 ± 6.28 | 72.2±5.51 | 74.75 ± 5.99 | $P=0.510$ |
| Male (%) | 55% | 55% | 87.5% | $P=0.229^+$ |
| MoCA<br>(mean ± sd) | 28.35 ± 1.14 | 27.55 ± 1.47 | 14.12 ± 5.14 | $P=0.000^*$ |
| BMI<br>(mean ± sd) | 25.75 ± 2.04 | 25.31±2.82 | 24.41 ± 3.20 | $P=0.532$ |
| Fasted total ghrelin<br>(pg/ml) (mean ± sem) | 541.0±87.98 | 647.9±108.7 | 377.8±77.00 | $P=0.2896$ |
| Fasted acyl-ghrelin<br>(pg/ml) (mean ± sem) | 78.93±13.77 | 114.4±18.82 | 51.39±17.24 | $P=0.0925$ |
| Post-prandial (180 min) total ghrelin<br>(pg/ml) (mean ± sem) | 500.4 ± 65.80 | 566.1 ± 96.89 | 401.8 ± 88.61 | $P=0.6464$ |
| Post-prandial (180 min) acyl-ghrelin<br>(pg/ml) (mean ± sem) | 72.86 ± 10.75 | 112.0 ± 18.31 | 41.32 ± 18.25 | $P=0.0169^*$ |

